# Genetic deletion of interleukin-15 is not associated with major structural changes following experimental post-traumatic knee osteoarthritis in rats

**DOI:** 10.1101/2021.04.15.440032

**Authors:** Ermina Hadzic, Garth Blackler, Holly Dupuis, Stephen J Renaud, C. Thomas Appleton, Frank Beier

## Abstract

Post-traumatic Osteoarthritis (PTOA) is a degenerative joint disease, leading to articular cartilage breakdown, osteophyte formation, and synovitis, caused by an initial joint trauma. Pro-inflammatory cytokines increase catabolic activity and may perpetuate inflammation following joint trauma. Interleukin-15 (IL-15), a pro-inflammatory cytokine, is increased in OA patients, although its roles in OA pathophysiology are not well characterized.

IL-15 levels appear to correlate to self-reported pain levels, and polymorphisms in the IL-15 receptor alpha gene correlate to a 1.5-fold increase in OA symptoms. This could be due to IL-15 effects on the activity of proteinases, such as matrix metalloproteinases (MMP) −1, −3, and −7. Here we utilized *Il15* deficient rats to examine the role of IL-15 in PTOA pathogenesis in an injury-induced model of OA. OA was surgically induced in *Il15* deficient rats and control wild-type rats to compare PTOA progression. Semi-quantitative scoring of the articular cartilage, subchondral bone, osteophyte size, and synovium was performed by two blinded observers. Analyses of articular cartilage damage, subchondral bone damage, and osteophyte formation revealed no significant difference between *Il15* deficient rats and wild-type rats following PTOA-induction. Similarly, synovitis scoring across 6 parameters found no significant difference between genetic variants. Overall, IL-15 does not appear to play a key role in the development of structural changes in this surgically-induced rat model of PTOA.

## Introduction

Osteoarthritis (OA) is a musculoskeletal disorder that presents a significant global burden, with over 303 million cases of hip and knee OA, which is expected to increase [1]. OA is a disorder of the entire joint, initially categorized by abnormal joint tissue metabolism and later by structural joint tissue changes. Each tissue reacts in its own way to the disease, as well as to the actions of the other joint tissues with changes including cartilage degeneration, subchondral bone remodeling, osteophyte formation, and synovial inflammation [2]. Post-traumatic osteoarthritis (PTOA) is a subtype of OA that develops after joint trauma, such as meniscal or ligament injury, and occurs most often in the lower extremities [3]. PTOA is estimated to account for 12% of all lower extremity OA cases, with costs of about $3 billion USD annually [4]. Although PTOA may affect any joint, knees and ankles are the most commonly affected joints [3].

A proposed mechanism of PTOA is the perpetuation of inflammation, in which some patients are unable to resolve the acute inflammation following joint injury and thus develop chronic inflammation with symptomatic PTOA [5]. Inflammation in arthritis is typically associated with rheumatoid arthritis (RA), a systemic disease with joint swelling and damage, although it plays a role in OA as well [6, 7]. OA inflammation is distinctly different from RA in many ways, with OA having lower levels of inflammatory proteins, less pronounced synovitis, and no response to conventional biologics used in RA, to name a few [7]. OA is characterized by a chronic, low grade form of inflammation that involves pro-inflammatory cytokine release, such as interleukin 1 (IL-1) and tumor necrosis factor alpha (TNFα) [7, 8]. Pro-inflammatory cytokines are present in healthy synovium, although there seems to be a higher presence of anti-inflammatory cytokines that successfully suppress inflammation, suggesting there is an imbalance in OA [9]. In fact, levels of IL-1*β* and TNF are increased in the synovial fluid, synovium, subchondral bone, and cartilage of the OA joint [10]. The prolonged increase in these pro-inflammatory cytokines within the joint leads to an increase in expression of downstream catabolic factors, such as matrix metalloproteinases (MMPs) [11]. These MMPs are zinc-dependent enzymes that play a major role in extracellular matrix (ECM) degradation through the cleavage of type II collagen and other ECM components (e.g., aggrecan, fibronectin, laminin) [12]. The inflammation of the synovium, termed synovitis, is an important aspect in this perpetuation of inflammation model, as the failure to resolve this inflammation results in drastic changes to the resident cells, leading to hyperplasia, angiogenesis, and infiltrates within the synovial membrane. Infiltrating cells, such as T-cells, B-cells, plasma cells, and macrophages, then contribute to further destruction of the cartilage matrix and subchondral bone [13].

Interleukin-15 (IL-15) is a pro-inflammatory cytokine that is crucial for natural killer cell (NK) ontogeny and CD8 T cell memory [14]. There are 3 receptors for IL-15: (1) the specific IL-15R*α*, (2) the shared IL-2/IL-15R*β*, and (3) the common IL-15R*γ*, binding IL-15, −2, −4, −7, and −19. IL-15 has a high affinity for the IL-15R*α* chain, but can only transduce signals in the presence of the IL-15*β* and *γ* receptors, via the JAK1/STAT3, JAK3/STAT5, and NF-*κ*B pathways [11, 14]. IL-15 is widely expressed by monocytes, macrophages, dendritic cells, fibroblasts, epithelial cells, and skeletal muscle [15]. Additionally, IL-15 signaling plays a vital role in the bone turnover process. IL-15R*α* ensures efficient osteoblast/osteoclast coupling, as well as determining osteoblast phosphate homeostasis and mineralization capacity [16]. Female *Il15Rα*^-/-^ mice have impaired osteoclast activity and are protected from age related trabecular bone loss when ovariectomized [17]. Indeed, single nucleotide polymorphisms (SNP) of IL-15R*α* positively correlate to total bone volume, as well as cortical bone volume [18]. Overall, IL-15 plays an important role in many body tissues and processes.

The relationship between IL-15 and RA has been more thoroughly investigated, although since OA-related inflammation is different from RA-related inflammation, potential roles of IL-15 in OA pathogenesis should be studied separately [19]. IL-15 protein and mRNA levels are reported to be significantly increased in patients with OA compared to the control group [20]. Further, Scanzello et al. (2009) found that IL-15 protein levels are higher in the synovial fluid during early OA compared to late stage, suggesting a role for IL-15 in the early stages of the disease [21]. Interestingly, another study found a positive correlation between SNPs in the human IL-15R*α* gene and prevalence of symptomatic OA [22]. Additionally, serum IL-15 levels independently correlate to pain intensity as measured by the Western Ontario McMaster University Osteoarthritis Index (WOMAC) pain score, although they do not correlate with Kellgren-Lawrence (KL) radiographic severity [23]. The increase of IL-15 in the synovial fluid of OA patients is also positively correlated to other pro-inflammatory cytokines, as well as MMP activity [24]. Specifically, Tao and colleagues (2015) demonstrated a strong correlation between MMP-7 serum levels and IL-15 [20]. A recent study showed that IL-15R*α* is present on chondrocytes, and treatment with IL-15 *in vitro* causes an increase in MMP1 and −3 release, but does not alter generation of soluble glycosaminoglycan (GAG) fragments [22]. Therefore, IL-15 may play a role in OA pathogenesis through an increase in pro-inflammatory activity that results in increased MMP activity. This MMP activity may then lead to increased tissue breakdown, and thus more pain and other symptoms.

In the present study, we investigated the role of IL-15 in the development of structural joint changes related to knee PTOA using a well-established rat model of surgically-induced PTOA in *Il15*^-/-^ rats and control *Il15*^+/+^ rats. We hypothesized that *Il15*^-/-^ rats would demonstrate a slower progression of PTOA compared to *Il15*^+/+^ rats.

## Materials and Methods

### Animals and surgery

Holtzman Sprague-Dawley rats possessing a 7 base pair frameshift deletion in the second exon of *Il15* were utilized, as described previously [25]. These rats do not produce mature IL-15 protein and were bred in-house. Rats were housed in a temperature- and humidity-controlled room (20-25 °C, 40-60%), with good ventilation. Water and standard rat chow were freely available. Rats were housed in colony cages and on a standard 12 hour light/dark cycle. Male *Il15*^+/+^ and *Il15*^-/-^ rats were then randomly allocated to either a surgical group (N = 15/genotype) or control (N = 9/genotype). The surgical group underwent anterior cruciate ligament transection with destabilizing medial meniscus (ACLT-DMM) surgery on the right knee at 9.5 weeks old, as described by Appleton et al. (2007) with one modification to reduce the speed of joint damage progression in this model [26]. Instead of performing partial medial meniscectomy, the medial meniscus was destabilized by transecting the medial meniscotibial ligament. Surgical anesthesia was induced using 5% Isofluorane, then decreased to 2% for maintenance. Ampicillin (40 mg/kg) was administered subcutaneously as a prophylactic antibiotic, and slow release Buprenorphine (1 mg/mL) was administered subcutaneously as a post-operative analgesic. A sham surgery is normally performed, with joint arthrotomy but no cruciate or meniscotibial ligament transection, although it was not used here as we did not want to induce any inflammation in the control group. Instead, we utilized age-matched surgically-naive rats as the control condition. Weights were measured daily for the first 4 days post-operatively, and then weekly. All rats were euthanized by asphyxiation with CO2 8 weeks after PTOA induction. All animal experiments were in accordance with the Canadian Council on Animal Care guidelines and were approved by the Animal Use Subcommittee at Western University (2019-029).

### Histopathology and Scoring

Right knees were dissected and fixed in 4% paraformaldehyde at 4 °C overnight. Decalcification with Decal Stat™ (StatLab, Baltimore, MD) and Formical-2000™ (StatLab, Baltimore, MD) was carried out on coronally bisected joints. Following processing and paraffin embedding, knees were frontally sectioned at 6 μm. Surface decalcification with Decal Stat™was used as required. The centre of the joint was analyzed by staining 3-5 serial sections, 200 μm apart, stained in 0.04% Toluidine Blue for cartilage and bone histopathology or with Hematoxylin and Eosin (H&E) for synovial histopathology analysis. Slides were randomized and blinded for scoring by 2 observers.

Toluidine Blue stained sections were analyzed using the Osteoarthritis Research Society International (OARSI) rat histopathologic system [27, 28]. Cartilage degeneration, subchondral bone damage, and osteophytes were graded in four knee quadrants - Medial Femoral Condyle (MFC), Medial Tibial Plateau (MTP), Lateral Femoral Condyle (LFC), and Lateral Tibial Plateau (LTP). For each animal, a score per parameter was assigned based on the mean, median, peak, and summed score for the slides. Within the summed score, there is a maximum score of 75 for total cartilage damage, 25 for subchondral bone damage, and 20 for osteophyte scores.

H&E-stained sections were analyzed using a six parameter synovial scoring system, assessing synovial lining thickness, sub-synovial infiltration, fibrin deposition, vascularization, fibrosis, and perivascular edema [29]. Scoring was assessed separately in the medial and lateral parapatellar, superior, and inferior compartments (6 compartments total). Each synovial histopathology parameter was assigned a score of 0 (none) to 3 (severe). Final scores for each synovial histopathology parameter were calculated for each animal by summation of the mean score (average of 3-5 slides per knee) from each of the 6 joint compartments, producing a maximum score of 18.

### Statistical Analysis

Statistical analyses were performed in GraphPad Prism v.8.2 and IBM SPSS Statistics v.23. SPSS was used for Cohen’s kappa and Chi-Squared analysis, and Prism for any remaining statistics. Cohen’s kappa was run for inter-rater reliability of cartilage degeneration, subchondral bone damage, and synovitis scoring. The Chi-Squared test of independence with Cramer’s V was run between groups for the presence of osteophytes. Normality was assessed via the D’Agostino & Pearson test. A two-way analysis of variance (ANOVA) with Tukey’s multiple comparison test was run for all OARSI histological scoring. Synovitis scores were analyzed using a one-way ANOVA with Tukey’s multiple comparison test, or the Kruskal-Wallis with Dunn’s multiple comparison test, depending on normality. Weight was compared using a Two-Way ANOVA with repeated measures, and Tukey’s multiple comparison test. P < .05 was considered statistically significant.

## Results

### No differences in weight gain between surgical groups or genotypes

In order to study PTOA in the rat model, OA was surgically induced at 9.5 weeks of age via ACLT-DMM surgery. Age-matched rats that did not have any kind of surgery were used as the controls, termed the Naive group. In order to assess whether surgery affected weight of *Il15*^+/+^ or *Il15*^-/-^ rats, all animals were weighed daily for the first 4 days post-operation (or timepoint start in the case of the Naive group), then weekly until the end of the 8 weeks. There was no significant weight difference between any groups at any timepoints (**Fig. 1**). The *Il15*^+/+^ PTOA rats were the only group to lose weight during the first 4 days post-op, although the change in weight in *Il15*^+/+^ PTOA rats was not significantly different compared to the remaining groups. Additionally, *Il15*^-/-^ rats, in both the PTOA and Naive groups, trend towards weighing less than the *Il15^+/+^*, although this was not statistically significant (*Il15*^+/+^ and *Il15*^-/-^ PTOA mean difference = 26.00, 95% CI [−14.59, 66.59] and Naive mean difference = 29.78, 95% CI [−32.42, 91.98]). The general health of all animals was deemed to be good, with no adverse events observed following surgery or otherwise.

**Figure 1 –.**
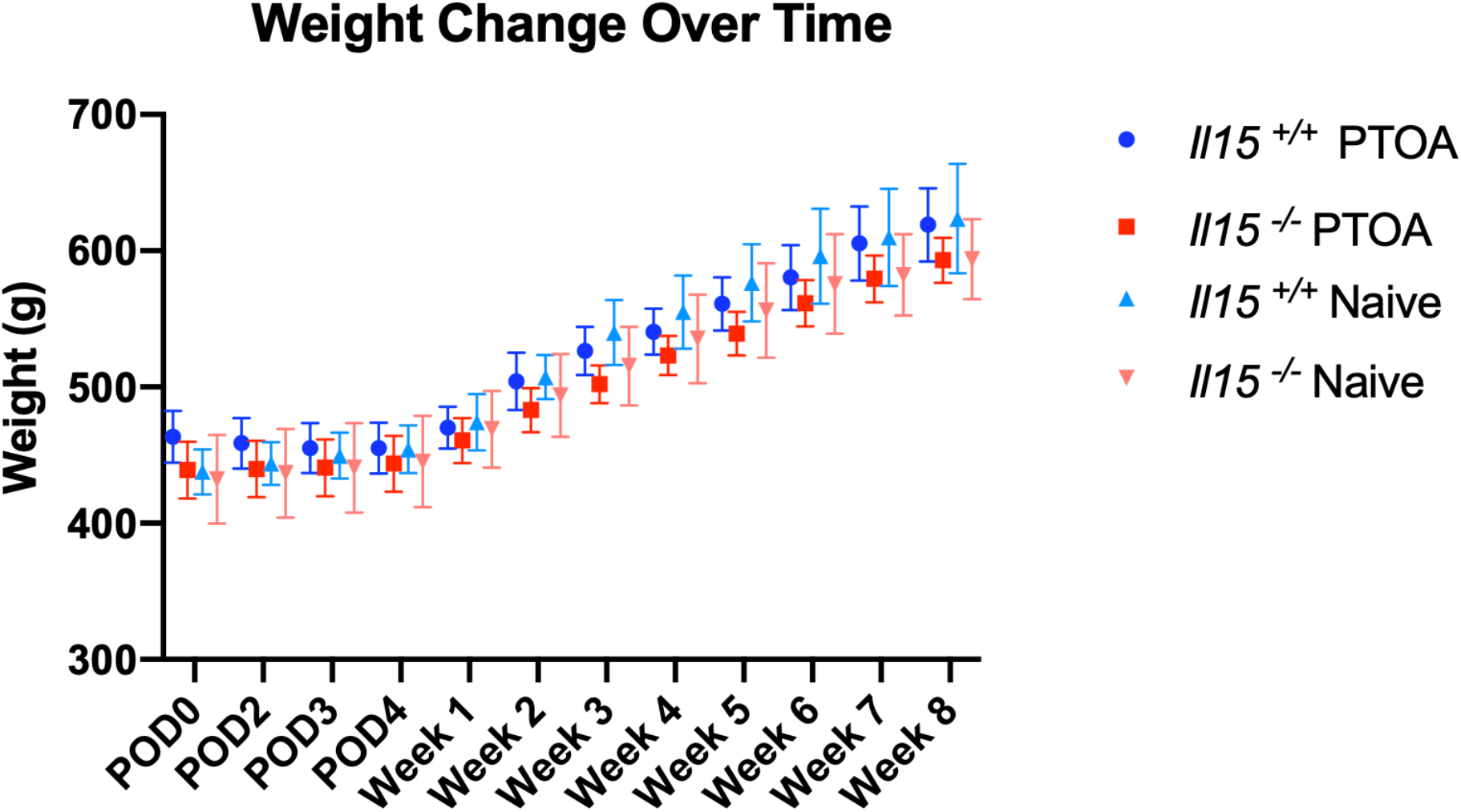
All groups gained weight similarly over the 8 weeks. Weight of animals, in grams (g), was measured daily for the first 4 days post-operation, and then weekly for 8 weeks. A Two-Way ANOVA with repeated measures, and Tukey’s multiple comparison was run post-hoc. There was no significant difference between the groups at any timepoint, with all groups steadily gaining weight throughout the experiment. (N = 15 rats/PTOA group, and N = 9 rats/Naive group, p > 0.05, data represented are mean with 95% CI).

### Cartilage damage after surgery is not affected by genotype

Following tissue collection, OARSI histological analysis was carried out by 2 scorers, using a semi-quantitative scoring system with a scale from 0 (no damage) to 5 (severe damage) to quantify articular cartilage damage across 4 quadrants. Further, the cartilage in each quadrant was assessed in 3 equal zones, with scores summed in order to represent the total cartilage damage (maximum total of 15). Reliability testing of cartilage histopathology scores revealed a weighted kappa score was 0.71, 95% CI [0.45, 0.51], demonstrating substantial rater agreement, according to the Landis & Koch (1977) guidelines [30]. The PTOA groups had evidence of cartilage damage (**Fig. 2**), but the Naive groups did not (**Fig. 3**). As expected, all PTOA groups had significantly higher OARSI scores compared to the Naive groups, which was more pronounced in the medial femoral and tibial cartilage, with the highest scores in the MTP (**Fig. 4**). However, damage across the articular cartilage was not significantly different between *Il15*^+/+^ PTOA and *Il15*^-/-^ PTOA rats in the mean, median, peak, and summed scores. The Naive group did not show consistent evidence of articular cartilage damage, which was consistent between genotypes. Zonal analysis revealed that Zone 2 (middle part of the plateau/condyle) had significantly higher scores for the PTOA group, although this was also non-significant when comparing *Il15*^+/+^ to *Il15*^-/-^ animals. Overall, while surgery induced clear structural changes in articular cartilage, damage does not appear to be significantly different between *Il15*^+/+^ PTOA and *Il15*^-/-^ PTOA rats.

**Figure 2 –.**
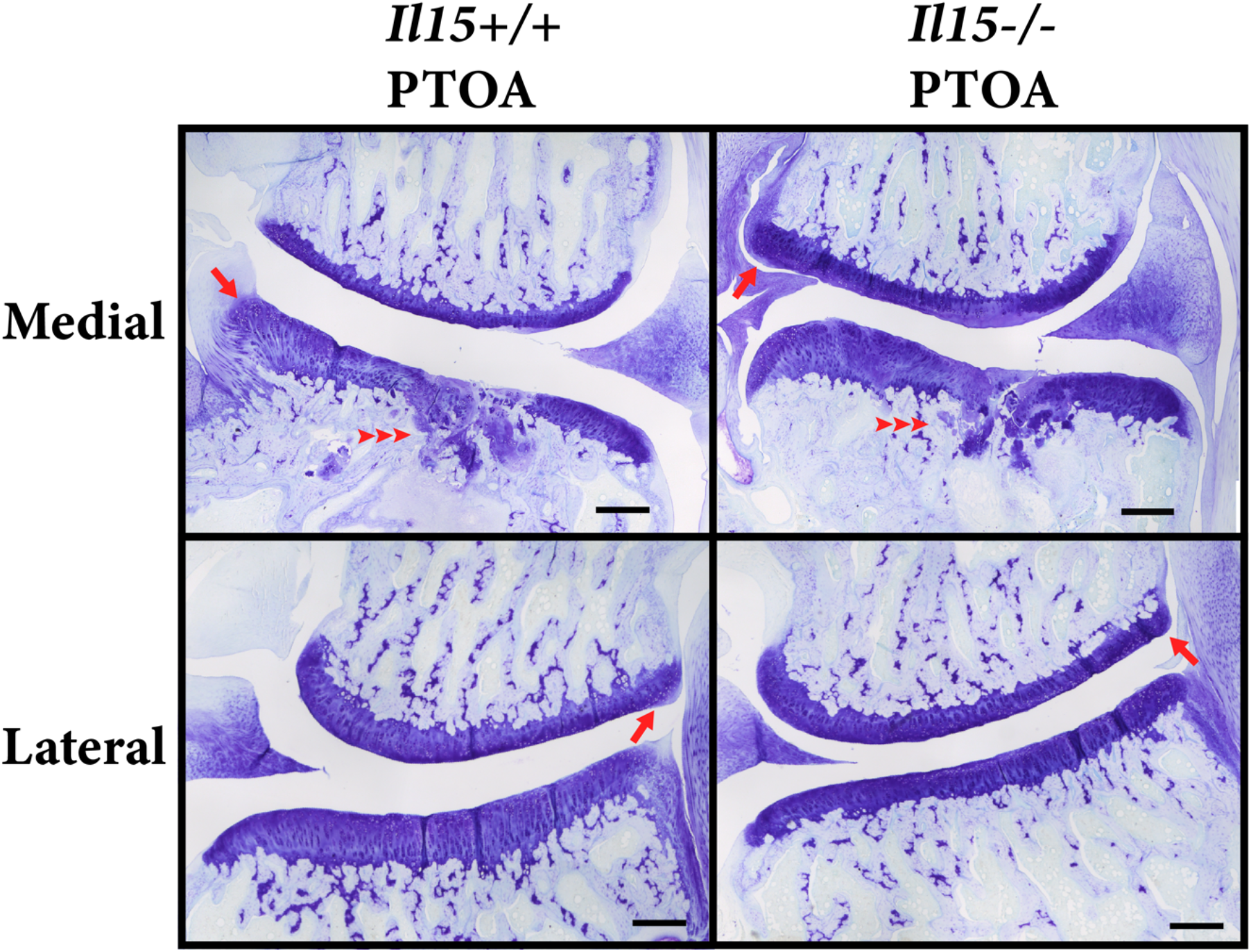
Representative histological images demonstrate greatest damage in the medial compartment of the PTOA group. Representative images of toluidine blue-stained paraffin sections of rat knees, demonstrating that the PTOA group had the greatest damage in the medial aspect as seen by osteophytes (red arrow), cartilage loss, and subchondral bone damage (red arrow heads). This damage was not significant different between the *Il15*^+/+^ PTOA and *Il15*^-/-^ PTOA rats. Images taken at 4X magnification, scale bars represent 200 μm (N = 15 rats/PTOA group).

**Figure 3 –.**
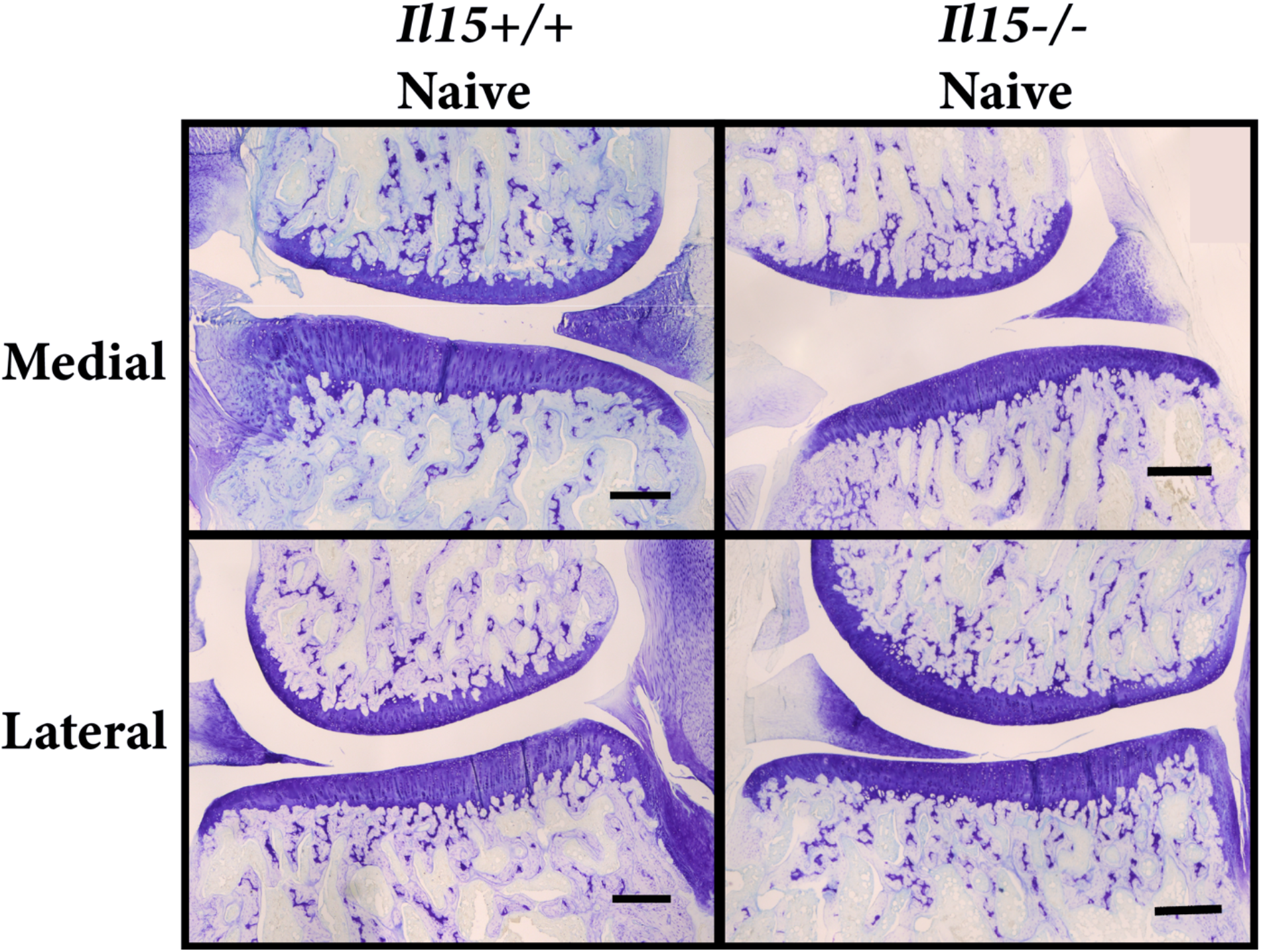
Representative histological images demonstrate healthy joint tissues in Naive rats. Representative images of toluidine blue-stained paraffin sections of rat knees, demonstrating that the Naive group had no significant differences between genotypes and was appeared healthy. Semi-quantitative scoring demonstrates no significant damage to the articular cartilage, subchondral bone, and osteophyte formation. Images were taken at 4X magnification, scale bars represent 200 μm (N = 9 rats/Naive group).

**Figure 4 –.**
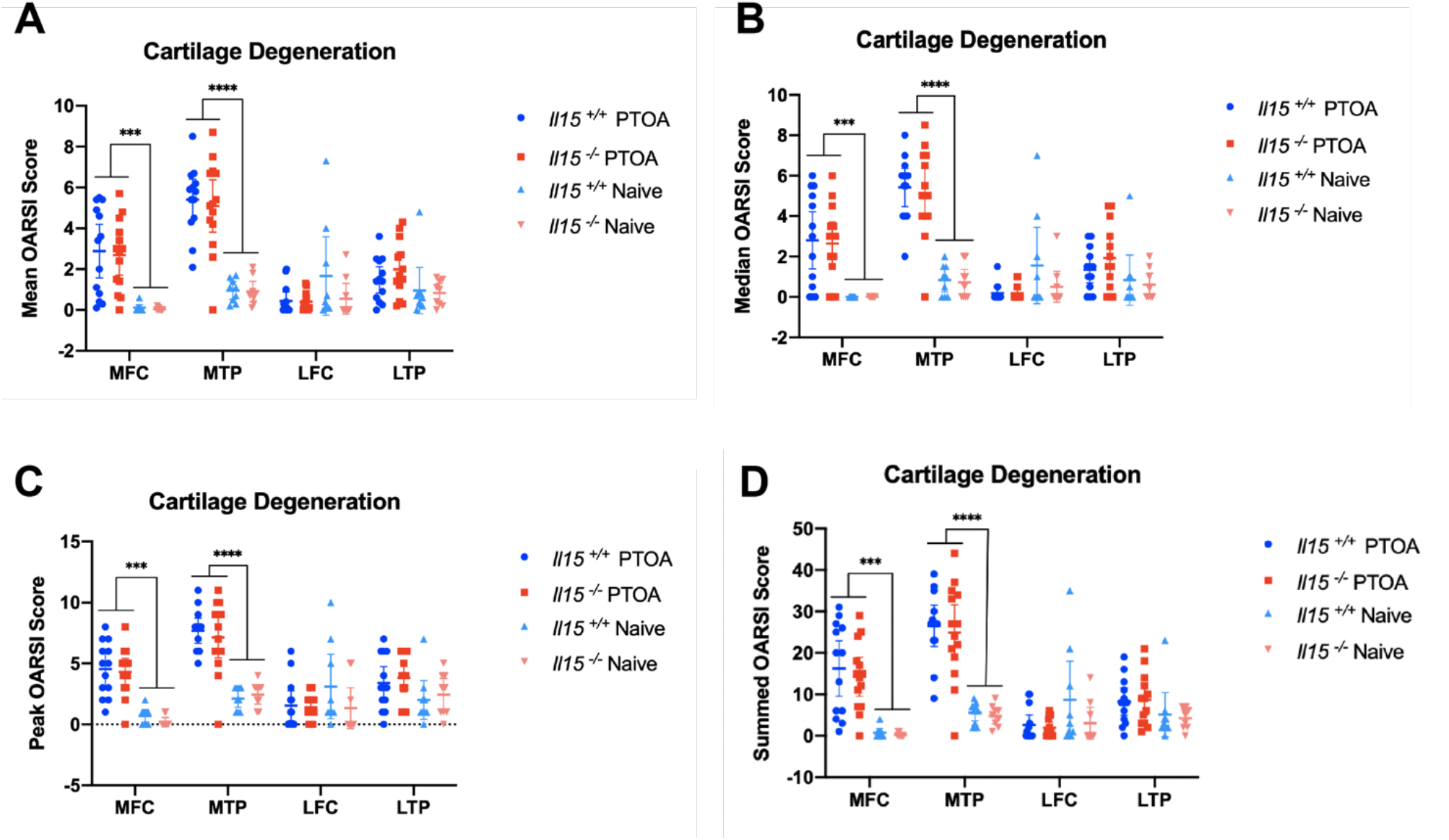
Cartilage damage is not significantly different between *Il15*^+/+^ PTOA and *Il15*^-/-^ PTOA animals. Animals either underwent ACLT-DMM surgery to induce PTOA or served as control with no surgery (Naive). Semi-quantitative scoring (scale 0 – 5) of toluidine blue-stained paraffin sections was used to determine the extent of cartilage damage in the joint via 4 compartments; medial femoral condyle (MFC), medial tibial plateau (MTP), lateral femoral condyle (LFC), and lateral tibial plateau (LTP). **A – D)** Two-Way ANOVA with Tukey’s multiple comparison test was used for statistical analysis; data represents mean with 95% CI. There was no significant difference in mean, median, peak, and summed scores between the *Il15*^+/+^ PTOA and *Il15*^-/-^ PTOA groups. Across all scores, the PTOA groups scored significantly higher than the Naive groups in the medial compartment. (*** p < 0.001, **** p < 0.0001, N = 15 rats/PTOA group and 9 rats/Naive group).

### Subchondral bone damage and osteophytes is similar between genotypes after surgery

Subchondral bone damage was assessed similarly to the articular cartilage damage, with a semi-quantitative score from 0 – 5. There was almost perfect inter-rater agreement in this parameter, with a weighted kappa of 0.86, 95% CI [0.60, 0.69], using the guidelines by Landis & Koch [30]. Only the medial tibial plateau demonstrated significantly higher scores in the PTOA groups compared to the Naive groups (**Fig. 5**). The mean, median, peak, and summed scores between the *Il15*^+/+^ PTOA and *Il15*^-/-^ PTOA rats was not significantly different in any of quadrants. The Naive group did not have consistent evidence of subchondral bone damage, which was consistent between genetic variants (**Fig. 3, 5**). Osteophytes were analyzed by recording the presence or absence by both raters, and subsequently measured by one rater. The presence of osteophytes was analyzed first, revealing a significant relationship between groups and osteophyte presence in the medial compartment (**Fig. 6.A**). Further analysis found this to be a strong association in both the medial femur and tibia, with the PTOA groups more likely to present with osteophytes (Cramer’s V = 0.64 and 0.60, respectively). In the lateral compartment, the same relationship was present in only the femoral condyle, with a moderate association (Cramer’s V = 0.43). Examination of the osteophyte size reveals no significant difference between the *Il15*^+/+^ PTOA and *Il15*^-/-^ PTOA rats in the mean, median, peak, and summed scores. Similarly, there is no significant difference within the Naive group (**Fig. 6.B-E**). Overall, there was no significant difference in the severity of subchondral bone damage or osteophytes between the *Il15*^+/+^ PTOA and *Il15*^-/-^ PTOA rats. Generally, OARSI scoring revealed greater OA damage in the medial compartment of the PTOA group (**Fig. 2**), with healthy tissue in the Naive group (**Fig. 3**).

**Figure 5 –.**
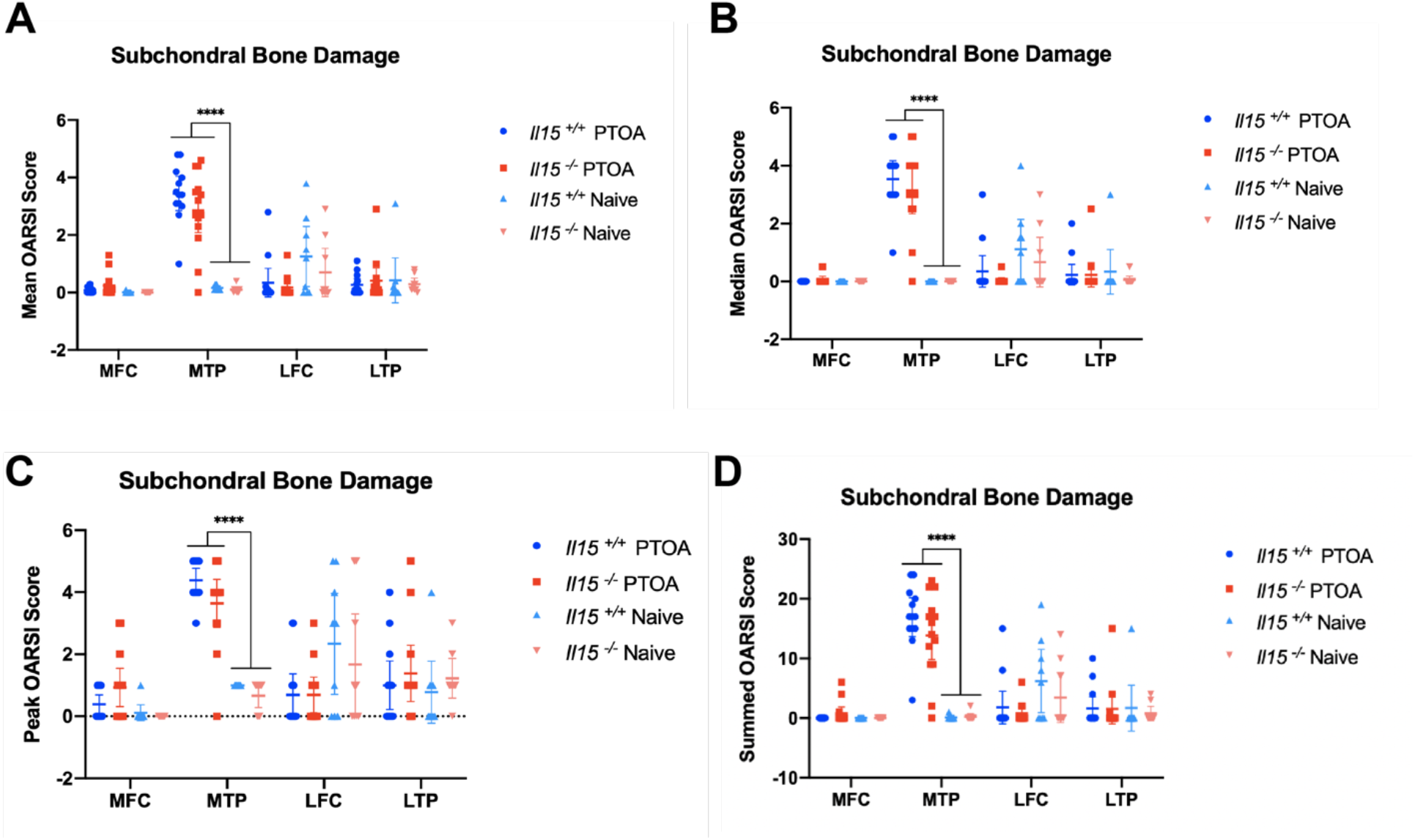
Subchondral bone damage is not significantly different between *Il15*^+/+^ PTOA and *Il15*^-/-^ PTOA rats. Animals either underwent ACLT-DMM surgery to induce PTOA or served as a control with no surgery (Naive). Semi-quantitative scoring (scale 0 – 5) was used to determine the extent of subchondral bone damage in the joint via 4 compartments; medial femoral condyle (MFC), medial tibial plateau (MTP), lateral femoral condyle (LFC), and lateral tibial plateau (LTP). **A – D)** Two-Way ANOVA with Tukey’s multiple comparison test was used for statistical analysis; data represents mean with 95% CI. There was no significant difference in mean, median, peak, and summed scores between the *Il15*^+/+^ PTOA and *Il15*^-/-^ PTOA groups. Across all scores, the PTOA groups scored significantly higher than the Naive groups only in the MTP. (**** p < 0.0001, N = 15 rats/PTOA group and 9 rats/Naive).

**Figure 6 –.**
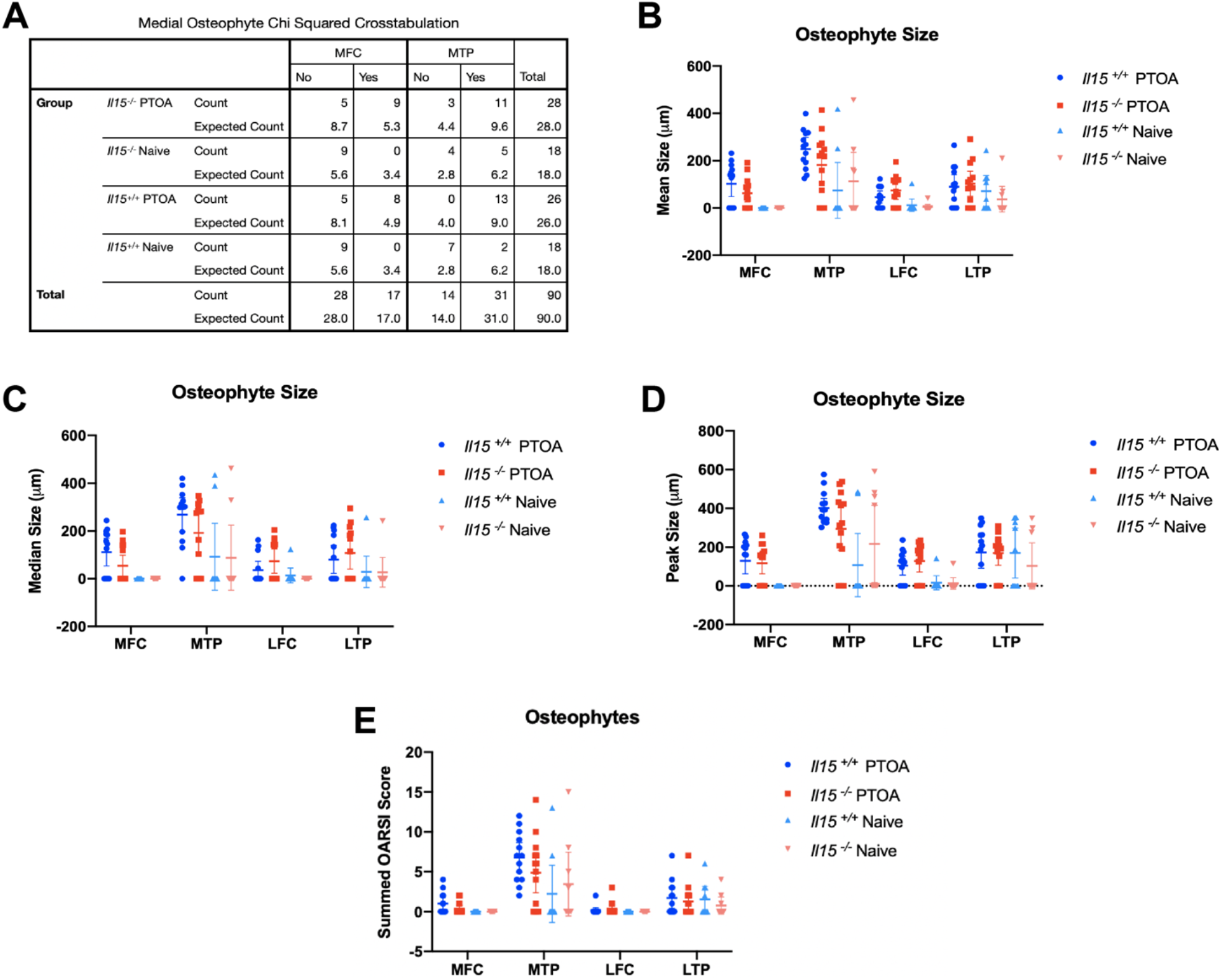
Osteophyte size is not significantly different between *Il15*^+/+^ and *Il15*^-/-^ animals. Animals either underwent ACLT-DMM surgery to induce PTOA or served as control with no surgery (Naive). Osteophytes were analyzed for absence/presence and size in the 4 compartments; medial femoral condyle (MFC), medial tibial plateau (MTP), lateral femoral condyle (LFC), and lateral tibial plateau (LTP). **A)** Presence of osteophytes was determined and agreed upon by two scorers. Chi-squared analysis with Cramer’s V established that the PTOA groups were strongly associated with osteophyte presence in the medial compartment, and moderately associated in the LFC, with no significance in the LTP. (MFC = X^2^ (3, N = 45) = 18.24, p < 0.0001, Cramer’s V = 0.64, MTP = X^2^ (3, N = 45) = 16.38, p < 0.005, Cramer’s V = 0.60, LFC = X^2^ (6, N = 45) = 16.90, p < 0.05, Cramer’s V = 0.43, LTP = X^2^ (3, N = 45) = 7.05, p > 0.05). **B – E)** Two-Way ANOVA with Tukey’s multiple comparison test was run, and data represents mean with 95% CI. There was no significant difference in mean, median, peak, and summed scores all demonstrate that there was no significant difference between the *Il15*^+/+^ PTOA and *Il15*^-/-^ PTOA groups. Mean, median, and peak represents osteophyte size, while the summed score represents the designated OARSI score (p > 0.05, N = 15 rats/PTOA group and 9 rats/Naive group).

### Synovitis is not affected by IL15 genotype

Synovitis was similarly assessed using a semi-quantitative 6 parameter scoring system. A score from 0 (none) to 3 (severe) was given across 6 compartments (medial and lateral parapatellar, superior, and inferior compartments) for 6 parameters; (1) synovial lining thickness, (2) sub-synovial infiltration, (3) surface fibrin deposition, (4) vascularization, (5) fibrosis, and (6) perivascular edema. Inter-rater reliability testing reveals substantial to almost perfect agreement, with a weighted kappa ranging from 0.67 to 0.85 across the six parameters [30]. As expected, signs of severe synovitis including vascularization, fibrosis, and vascular edema were similar between PTOA and Naive groups at this early stage of PTOA development. In keeping with early stage PTOA development, only the sub-synovial infiltrate and surface fibrin scores were increased in PTOA animals. However, there was no clear difference between genetic strains (**Fig. 7**). There was no significant statistical difference between the *Il15*^+/+^ and *Il15*^-/-^ rats across all six parameters (**Fig. 8**). Overall, synovitis did not clearly differ between the *Il15*^+/+^ PTOA and *Il15*^-/-^ PTOA rats 8 weeks after PTOA induction.

**Figure 7 –.**
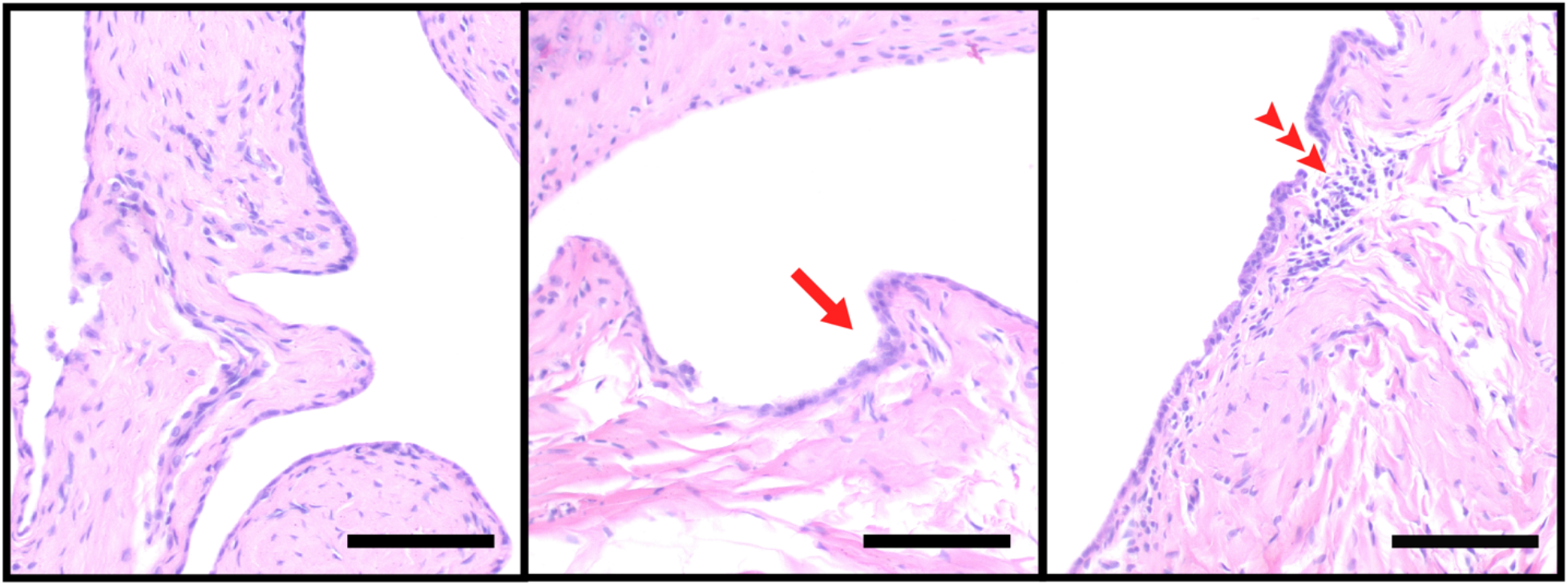
Representative images of synovitis in the PTOA group. Histological images of the synovitis, as demonstrated by surface fibrin and sub-synovial infiltration, found in the PTOA animals. Healthy synovium (first box) is presented as a comparison and represents Naive tissue. Surface fibrin (red arrow) and sub-synovial infiltration (red arrowheads) was found significantly more in the PTOA group compared to the Naive, but was not different between genetic variants. Images take at 20X magnification, scale bars represent 100 μm (N = 15 rats/PTOA group and N = 9 rats/Naive group).

**Figure 8 –.**
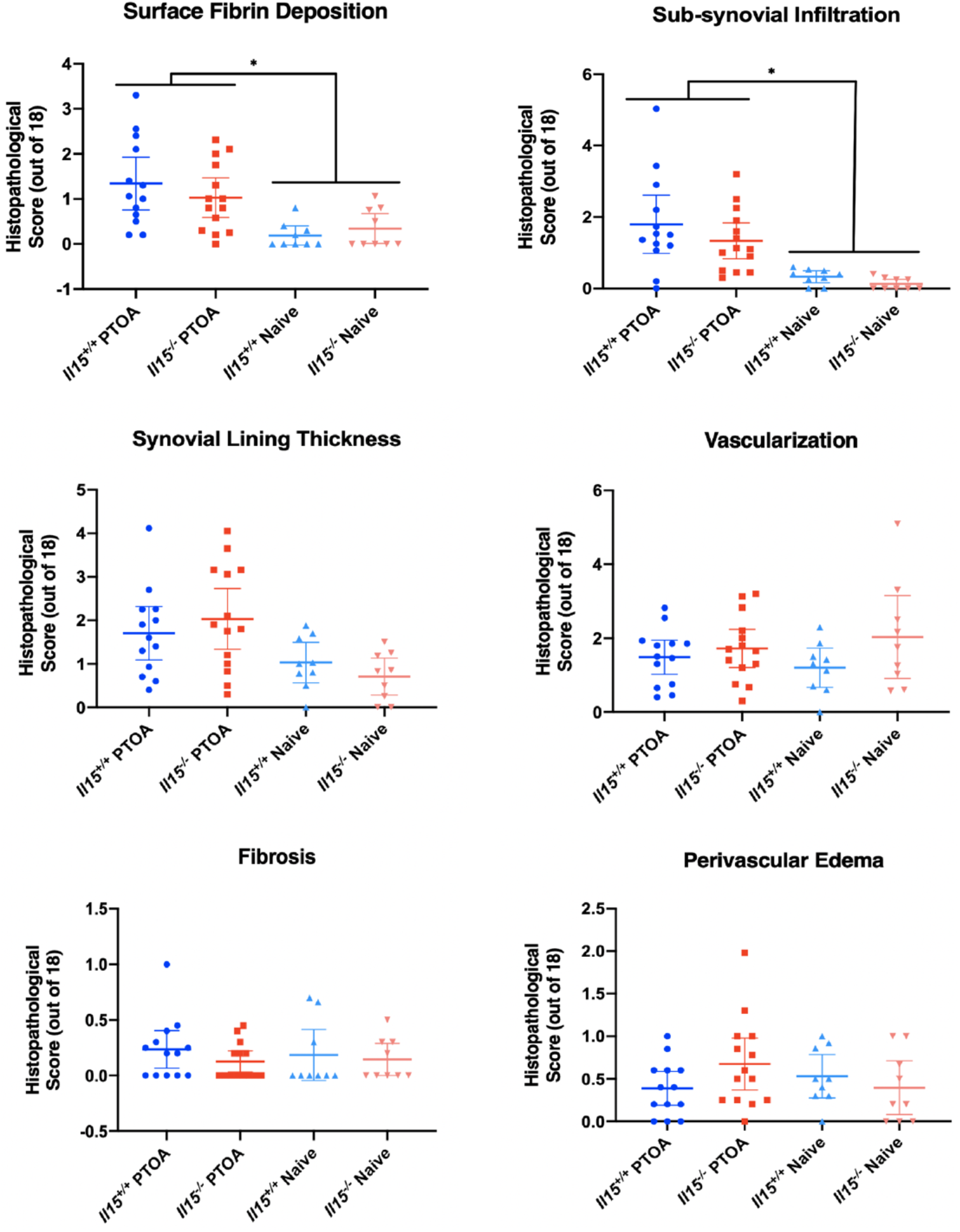
Synovitis is not significantly different between *Il15*^+/+^ and *Il15*^-/-^ animals. Rats either underwent ACLT-DMM surgery to induce PTOA or served as control with no surgery (Naive). Semi-quantitative scoring was used to assess synovitis across 6 parameters, ranging from 0 (none) to 3 (severe), measured in 6 zones (medial and lateral parapatellar, superior, and inferior compartments). A One-Way ANOVA with Tukey’s multiple comparison test or the Kruskal-Wallis with Dunn’s multiple comparison test was run, depending on normality. Data represents mean with 95% CI. Analysis reveals that the PTOA group had scored significantly higher for sub-synovial infiltration and surface fibrin, although there was no significant difference between genetic strains. The remaining 4 parameters were non-significant across all groups (* p < 0.05, N = 15 rats/PTOA group and 9 rats/Naive group).

## Discussion

Cytokine activity is a well-defined research area in OA, offering many potential therapeutic targets to be explored. Interleukin-15 (IL-15), a pro-inflammatory cytokine, with important roles in innate immunity, has increased levels in OA tissue and can potentially increase catabolic activity within the joint. Our study utilized a Holtzmann Sprague-Dawley male global *Il15*^-/-^ model to study the contribution of IL-15 in PTOA progression. Our work demonstrated that *Il15*^-/-^ rats present with normal joint morphology in the absence of surgery and display similar structural features of PTOA to their *Il15*^+/+^ counterparts. Through the analysis of cartilage damage, subchondral bone damage, osteophyte formation, and synovitis we did not see any significant differences between genotypes, suggesting that IL-15 does not play a vital role in PTOA progression in this rat model, at least at the stage investigated. These data do not support our initial hypothesis.

Weight was recorded as a measure of general health, especially for the PTOA group following surgery. There was no significant difference in weight at any timepoint between all groups, similar to the findings from Renaud et al. (2017) who compared weight of *Il15*^+/+^ and *Il15*^-/-^ female rats [25]. Even though the *Il15*^+/+^ PTOA rats lost weight during the first 4 days post-op, this loss was not statistically significant, and they continued to gain weight in the weeks following. Interestingly, the *Il15*^-/-^ rats, in both the PTOA and Naive groups, consistently weighed less than their *Il15*^+/+^ counterparts, although not enough to reach statistical significance. *Il15*^-/-^ mice have varied in their reported weight differences in the literature; for example, Kennedy et al. (2000) reported that *Il15*^-/-^ mice were not significantly different in body weight compared to *Il15*^+/+^ mice, but another study reported that female *Il15*^-/-^ mice weighed less [31, 32]. Similarly, studies using *Il15Rα*^*-/-*^ mice have conflicting reports of weight, either reporting similar weights or a decrease in weight in mutants compared to controls [33, 34]. Interestingly, *Il15*^-/-^ mice on a high fat diet are resistant to diet-induced weight gain due to increased thermogenic capacity in brown and beige fat cells [35]. Our male *Il15*^-/-^ rats trend towards weighing less, perhaps due to a similar increased thermogenic capacity, although not enough to reach statistical significance at the timepoints used in our study.

In order to assess PTOA progression, ACLT-DMM surgery was utilized in both *Il15*^-/-^ and *Il15*^+/+^ rats. Early stage PTOA was successfully induced 8 weeks following surgery, as determined by statistically higher scores for cartilage damage, subchondral bone damage, and osteophyte presence compared to the Naive group. As expected, the damage was generally contained to the more weight bearing medial compartment. There is also evidence of synovitis developing in the PTOA group, as they scored significantly higher for sub-synovial infiltration and surface fibrin deposition, compared to the Naive group. Overall, ACLT-DMM surgery successfully induced mid-stage PTOA.

Inconsistent with our hypothesis, there was no significant difference between the *Il15*^-/-^ and *Il15*^+/+^ PTOA rats, indicating that IL-15 may not play a vital role in rat PTOA pathophysiology. This finding is also inconsistent with previous studies that found a relationship between IL-15 activity and OA, which may be due to a few factors. Firstly, it is possible that the rodent model is not ideal for translating studies in IL-15, as current literature on IL-15 and OA has exclusively been reported from human studies [20–24]. Additionally, our study examined only the 8-week timepoint, but the activity of IL-15 may be more robust at an earlier or end stage of disease. Another pitfall could be due to differing OA phenotypes. The current literature examining IL-15 and OA has excluded patients with a history of traumatic joint injury [20–24], thus excluding PTOA, which could be another reason for the lack of significance between groups in our study. IL-15 may play a more vital role in other OA phenotypes, such as primary OA, where risk factors such as ageing and metabolic syndrome may have effects on OA pathophysiology that involve IL-15, but were not examined in this study. Research utilizing *Il15*^-/-^ mice on a high fat diet revealed an important role for IL-15 in metabolic syndrome, as it seems to promote chronic inflammation in adipose tissue [35]. Perhaps the inflammatory role of IL-15 is more substantial in metabolic OA than in PTOA. Finally, the redundant and pleiotropic nature of cytokines could explain why the absence of IL-15 did not cause a change in PTOA progression. Additionally, the activity of IL-15 on catabolic factors, like MMPs, may be more indirect, as suggested by recent work by Warner and colleagues (2020). Chondrocytes treated with IL-15 *in vitro* demonstrate a delayed release of MMP-1 and −3, compared to TNF*α*, which may point to an indirect effect of IL-15 on these MMPs [22]. Therefore, the action of IL-15 to increase catabolic activity in the joint may be taken over by another cytokine when IL-15 is absent, especially if this activity is indirect.

It is important to consider the limitations involved in this work. Many of these limitations were due to the laboratory shutdown during the COVID-19 pandemic, which intervened with completion of additional *in vivo* experiments. Future work would benefit from an optimized method of IL-15 detection in the joint through immunohistochemistry in order to assess the expression of IL-15 in the rodent joint. As previously discussed, exploring multiple timepoints and perhaps a metabolic or aging OA model could be beneficial to exploring the role of IL-15 in OA. Pain and behavioral responses could not be fully assessed in our model due to restrictions; although it seems plausible that IL-15-deficiency could result in reduced pain despite normal structural progression of OA. Future studies should therefore include an examination of pain in this model. In addition, we were only able to study male rats, so inclusion of female rats in future studies would be beneficial.

Further, the use of a solely *in vivo* model limits the scope of our work in understanding the mechanisms of IL-15 in a more complex manner. An *in vitro* model studying primary rat or human synoviocytes that are treated with IL-15 would be of interest, investigating the effects and mechanism of IL-15 in the synovium. Finally, our ACLT-DMM model is invasive and does not perfectly mimic the injuries associated with PTOA in patients. Future work could utilize a non-invasive model, where the ACL is ruptured via tibial compression in order to induce PTOA [36, 37].

Overall, while our work did not demonstrate a significant difference in IL-15 and PTOA, there are still many avenues to explore with IL-15 to elucidate its potential role in OA pathogenesis.

